# Direct RNA targeted transcriptomic profiling in tissue using Hybridization-based RNA In Situ Sequencing (HybRISS)

**DOI:** 10.1101/2020.12.02.408781

**Authors:** Hower Lee, Sergio Marco Salas, Daniel Gyllborg, Mats Nilsson

## Abstract

Highly multiplexed spatial mapping of multiple transcripts within tissues allows for investigation of the transcriptomic and cellular diversity of mammalian organs previously unseen. Here we explore the possibilities of a direct RNA (dRNA) detection approach incorporating the use of padlock probes and rolling circle amplification in combination with hybridization-based *in situ* sequencing (HybISS) chemistry. We benchmark a dRNA targeting kit that circumvents the standard reverse transcription limiting, cDNA-based *in situ* sequencing (ISS). We found a five-fold increase in transcript detection efficiency when compared to cDNA-based ISS and also validated its multiplexing capability by targeting a curated panel of 50 genes from previous publications on mouse brain sections, leading to additional data interpretation such as *de novo* cell typing. With this increased efficiency, we maintain specificity, multiplexing capabilities and ease of implementation. Overall, the dRNA chemistry shows significant improvements in target detection efficiency, closing the gap between the gold standard of fluorescent *in situ* hybridization (FISH) based technologies and opens up possibilities to explore new biological questions previously not possible with cDNA-based ISS, nor with FISH.

## INTRODUCTION

There are a wide array of technologies for *in situ* visualization of transcripts having various benefits and drawbacks^(1–3)^ with many current methods requiring specialized microscopes to resolve diffraction limited spots^(4–7)^. Although many fluorescent *in situ* hybridization (FISH)-based methods can have very high detection efficiency and/or high multiplexing capabilities, the upscaling of these technologies are not suited for cell typing projects such as the Human Cell Atlas^(8)^ or large cohort studies. Wrangling these datasets across large areas and samples is essential to gain medically relevant knowledge where individual samples don’t suffice.

Our lab has developed *in situ* sequencing (ISS) as a method for multiplex detecting transcripts within tissue based on barcoded padlock probes (PLPs) and rolling circle amplification (RCA), forming single stranded DNA repeats known as rolling circle products (RCPs)^(9,10)^. The latest iteration of ISS, hybridization-based *in situ* sequencing (HybISS)^(9)^, RCPs contain combinatorial barcodes that can be decoded by hybridizing primary probes then fluorescently labelled oligonucleotides over multiple cycles and visualized using conventional widefield fluorescence microscopes. Although ISS has been shown to have good signal detection and throughput, it suffers from low transcript detection efficiency^(10,11)^ that can be mainly attributed to the inefficiency of cDNA synthesis which has been used to improve the specificity for hybridization and ligation of PLPs. Direct hybridization and probing of mRNA *in situ* for improved efficiency has been attempted with different commercially available ligases, including Chlorella virus DNA ligase (PBCV-1 DNA ligase), but showed high tolerances of mismatches for ligation^(12)^, and consequently worse specificity compared to cDNA template DNA ligation using thermophilic DNA ligases. Recently, it was reported that T4-RNA Ligase 2 showed good ligation efficiency with 3’-RNA/5’-DNA PLPs (Chimeric PLPs) on RNA templates, and exhibits higher ligation fidelity on single nucleotide variations, compared to PBCV-DNA ligase^(13)^. Here we evaluate such a direct RNA (dRNA) chemistry *in situ* with HybISS (HybRISS: Hybridization-based <u>RNA</u> *in situ* sequencing) that targets RNA with chimeric PLPs while still retaining the fundamental benefits of ISS technology.

Combined with sequence-by-hybridization detection chemistry of HybISS, we applied a targeted gene panel on mouse coronal brain sections for a comparative analysis of methods and demonstration of its capabilities and potential. We show over five-fold increase in transcript detection efficiency when compared to cDNA-based ISS, leading to additional data interpretation such as *de novo* cell typing. With this increased efficiency, we maintain specificity, multiplexing capabilities and ease of implementation. Overall, a dRNA-based approach closes the gap between the gold standard of FISH-based detection efficiency while maintaining a high level of multiplexing and throughput.

## RESULTS

### Increased targeting efficiency and retained specificity of dRNA-HybRISS

The HybRISS method bypasses cDNA synthesis step using gene specific chimeric PLPs, before they are ligated, amplified by RCA and fluorescently labelled for detection (**Fig. 1A**). In order to make use of HybISS detection chemistry^(9)^, customized PLP backbone sequences contain 20 nucleotide (nt) long unique ID sequences that are assigned to each gene of interest to be decoded in a combinatorial manner by first binding ID sequence specific bridge probes that are then template for fluorophore conjugated detection oligonucleotides (DOs) (**Fig. S1A and Table S1**).

**Figure 1:**
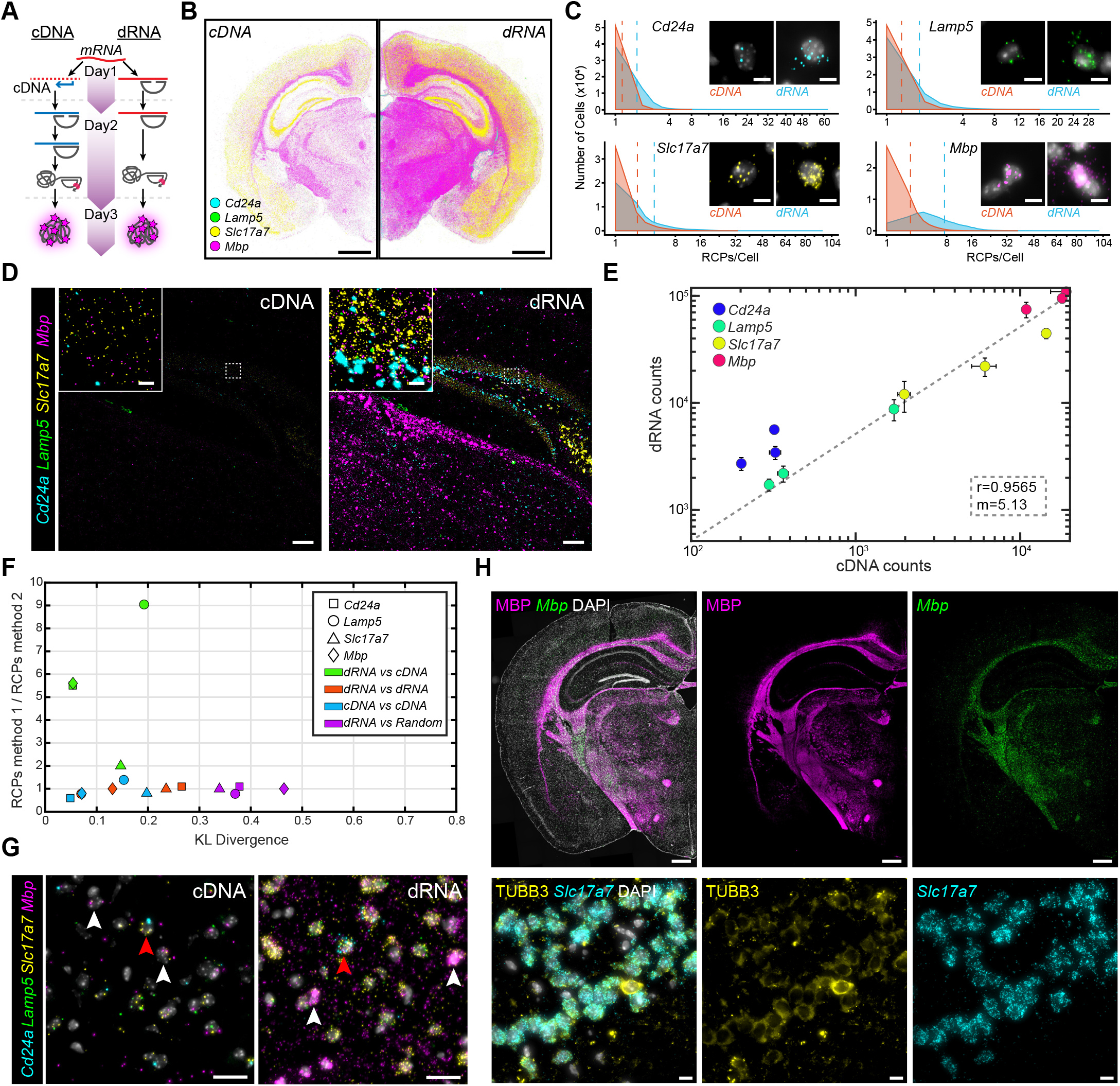
HybRISS: dRNA targeting in situ using a 4-plex gene panel. (**A**) Schematic of benchmarking experiment to compare cDNA- and dRNA-based HybISS. (**B**) Expression distribution of 4-plex gene panel (Cd24a/Lamp5/Slc17a7/Mbp) across sequential half coronal mouse brain sections. Displayed as output from MATLAB analysis pipeline. Scale bar, 1 mm. (**C**) RCP counts per cell of half coronal section and their frequency for each gene in the 4-plex panel. Representative raw images of single cells. Dashed lines represent the mean RCPs/Cell count for the respective chemistries. Scale bar, 5 μm. (**D**) Representative raw image of 4-plex from one of three ROIs (Fig. S2B,C). Experiments run in parallel and same postprocessing intensity level adjustments. ROIs include regions of cortex, hippocampus, and lateral ventricle. Scale bar, 100 μm, inset 10 μm. (**E**) Correlation plot of total RCP counts of dRNA against cDNA in three ROIs. x-axis represents cDNA and y-axis dRNA. Each spot of same color represents the three ROIs and deviation calculated from consecutive sections. (**F**) Kullback-Leibler divergence plot for the spatial distribution of 4-plex genes across a cortical ROI between cDNA and dRNA. x-axis represents spatial differences and y-axis represents frequency differences. (**G**) Multiplexed distribution of 4-plex genes in cortical region. Cd24a^+^ cells indicated by red arrowheads, Mbp^+^ cells indicated by white arrowheads. Scale bar, 20 μm. (**H**) Colocalization of fluorescent immunohistochemistry with dRNA-HybRISS. Top panels show MBP protein detection with Mbp dRNA-HybRISS. Bottom panels show pan-neuronal marker TUBB3 with excitatory neuron marker Slc17a7. Scale bar, top 500 μm, bottom 10 μm.

We first compared dRNA-HybRISS to cDNA-HybISS by targeting four genes (*i.e.* 4-plex) selected for their specificity in marking different cell types including ependymal (*Cd24a*), oligodendrocytes (*Mbp*), and excitatory neurons (*Lamp5* and *Slc17a7*) (**Fig. 1A,B and Fig. S1B**). PLPs were designed to target complementary sequences and the four genes could be discriminated from each other in a single cycle, but the possibility of combinatorial decoding is still feasible (**Fig. S1C**). To get an overall impression of the increased efficiency of dRNA, the total number of RCPs counted per segmented cell in the sections showed an overall increase in number and frequency for all four genes (**Fig. 1C**). This is visually clear when comparing images of single cells expressing the various genes (**Fig. 1C** insets, **Fig. S2A**). Furthermore, we selected three regions of interest (ROIs) encompassing the cortex, hippocampus and lateral ventricle for more detailed analysis (**Fig. S2B**).

Comparable images of the ROIs showed clear increased detection efficiency where sub-regional localization of detection could be seen with various densities in the dRNA condition, clear *Mbp* abundance in the corpus callosum, *Cd24a* surrounding the ventricle and *Slc17a7* within the cortex (**Fig. 1D and Fig. S2C**). Total sum of RCPs for each gene was quantified in the ROIs in replicate sections for each condition and found a correlation with a slope of 5.13, indicating an over five-fold increase in detection efficiency using the dRNA approach (**Fig. 1E**). Comparing RCP intensity showed variable results depending on the fluorophore (**Fig. S2D,E**). Overall, the signal-to-noise ratio (SNR) of dRNA-HybRISS were comparable to cDNA-HybISS, showing that RCA was not impeded in dRNA-HybRISS (**Fig. S2F,G**), consistent with the observation that the phi29 DNA polymerase used in RCA is as efficient on RNA/DNA chimeric circles as DNA circles^(14)^.

The increased number of detection events could be in part a consequence of off-target detection events. To evaluate this, we looked into the spatial distribution of the four targeted genes. Due to the architectural organization of the cortex, we were able to assess the spatial distribution of the four genes with a Kullback-Leibler divergence analysis along the cortex, observing a change in expression (y-axis) but no significant difference in spatial distribution (x-axis) between dRNA and cDNA (**Fig. 1F**). We further looked into ROIs to observe the spatial distribution of the four genes in more detail. Although cells expressing certain genes could be found within the cortex with both methods (**Fig. 1G**), it was visually clearer in dRNA approach as indicated by *Mbp* (white arrowheads) and *Cd24a* (red arrowhead) expressing cells. Within the lateral ventricle (**Fig. S3A**, top), a clear delineation of *Cd24a*^+^ cells lining the ventricle with dRNA, whereas it is sparser in cDNA. Furthermore, within the hippocampal formation, a clear separation of *Cd24a*^+^ and *Slc17a7*^+^ cells could be seen that was almost indistinguishable in cDNA approach (**Fig. S3A**, bottom). Co-localization of protein detection and RNA expression by performing immunohistochemistry (IHC) alongside dRNA-HybRISS revealed near identical staining pattern and density across the mouse brain tissue section when comparing *Mbp* (**Fig. 1H**, top). When targeting *Slc17a7* together with pan-neuronal marker Tubulin Beta-III (TUBB3) antibody (**Fig. 1H**, bottom), *Slc17a7* expression co-localized with most of the cells detected with TUBB3 within an ROI expressing *Slc17a7*^+^ cells and no RCPs were observed in cells that were not TUBB3 detected. Comparing the 4-plex dRNA approach to the Allen Mouse Brain Atlas in a cortical ROI, we see overlapping distribution of expression of all genes (**Fig. S3B**). This also applied to other regions as well as overall distribution of expression in the entire coronal section (**Fig. S3C**).

Control experiments to evaluate unspecific binding of the dRNA probes was done by switching 4-plex probe sets for the different experimental setups, but added an additional set of cDNA reference probes (*Actb*, *Gapdh*, *Pgk1*, and *Polr2a*) into the mix of dRNA probes for cDNA protocol (**Fig. S4A**). As expected, no signal was observed in either condition, only after stripping and labelling with bridge probes for the reference probes, RCPs could be visualized (**Fig. S4B**). A competitive assay using primers of varying concentration targeting the *Mbp* binding sites for the PLPs (**Fig. S4C**) resulted in almost a complete suppression of detectable *Mbp* RCPs (**Fig. S4D**).

### Multiplexing capacity of dRNA-HybRISS for de novo cell typing in mouse brain sections

To test the application and potential of the dRNA-HybRISS, we targeted a panel of 50 genes (*i.e.* 50-plex) curated based on previous publications to map cortical and hippocampal cell types: 33 genes from Codeluppi *et al.*^(4)^ and 17 from Qian *et al.*^(11)^ (**Fig. S5A, and Table S1**). Targets were probed sequentially over 14 rounds and then merged to create a composite image (**Fig. S5B and S6A-D**). The expression map obtained was then segmented to cells based on nuclear DAPI staining. Due to the increased RCP count per cell, we could perform *de novo* clustering on the data to resolve 56 clusters (**Fig. 2A,B and Supplementary Note 1**). While most of the neuronal cell types do not present unique markers in the panel, non-neuronal clusters were easily characterized by the expression of cell-type specific markers, whereas both excitatory and inhibitory clusters had a similar expression pattern between them, resulting in difficulties for discriminating analogous cell types within excitatory and inhibitory cells. These cell clusters can then be mapped back to a spatial position in the tissue for further analysis (**Fig. 2C**).

**Figure 2:**
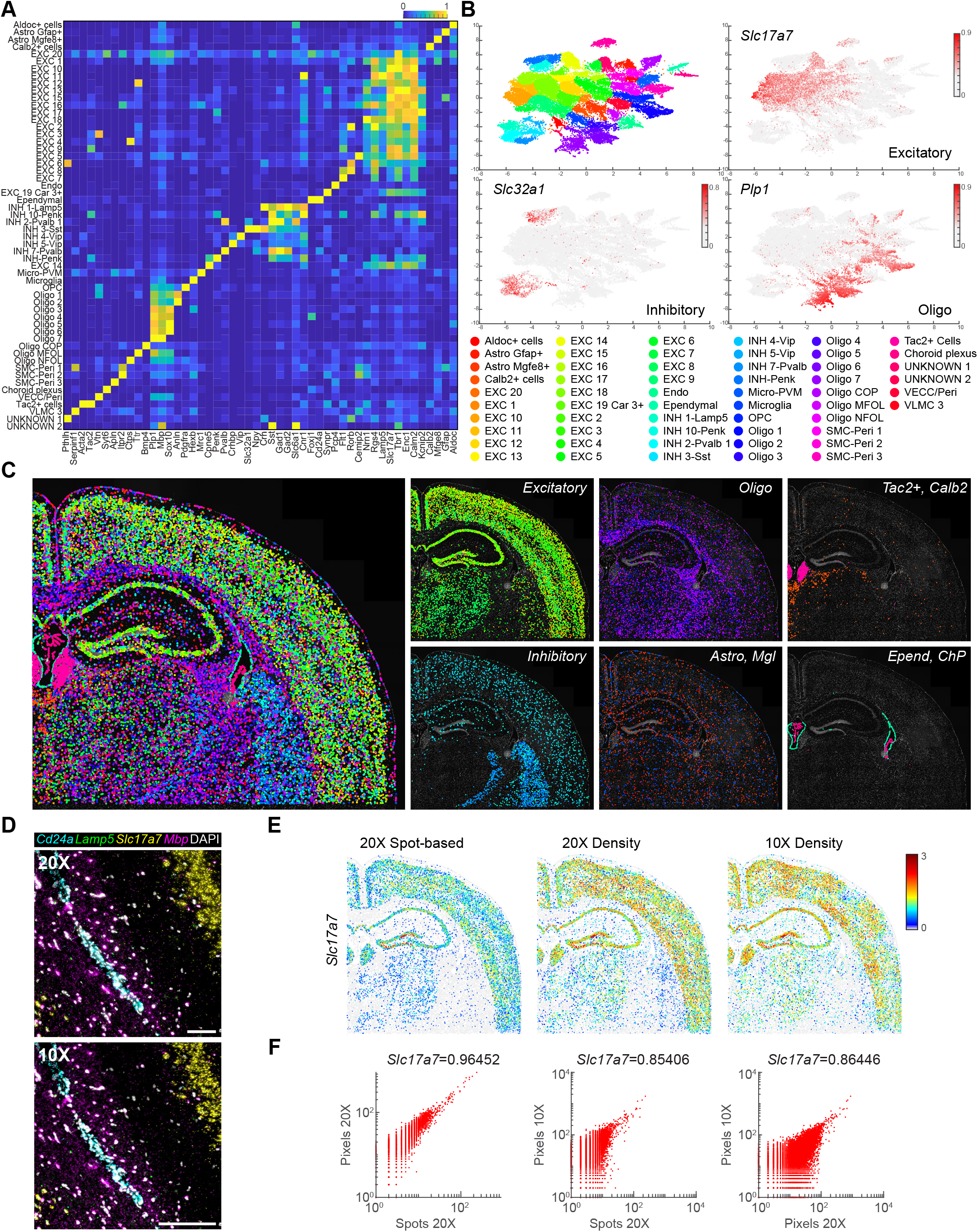
*De novo* clustering of 50 gene expression in mouse brain coronal section. (**A**) Expression matrix of 50 targeted genes across annotated cell clusters in segmented cells of the imaged region. (**B**) UMAP with de novo cell clustering based on the expression profile of the 50-plex gene panel. Three genes highlighted for their expression to mark pan-excitatory neurons (Slc17a7), inhibitory neurons (Slc32a1), and oligodendrocytes (Plp1). (**C**) Cell-type map across mouse coronal section, highlighting some classes in right panels. (**D**) Raw image comparison of 20X and 10X objective imaging. 200 pixel scale bar, 20X=64.2 μm, 10X=128.4 μm. (**E**) 20X objective spot-based detection converted to density-based detection compared to 10X objective density-base detection for Slc17a7. (**F**) Correlation comparison of 20X spot- and density-based detection to 10X-density based detection.

Additionally, due to amplified signals allowing for rapid imaging and now increased detection efficiency with dRNA, imaging at lower magnification to further increase throughput is possible. In parallel, we obtained a dataset with 10X imaging as a proof of concept (**Fig. 2D**). Here we produced expression maps based on a density threshold and assigned them to segmented cells (**Fig. 2E and Fig. S7A**). Indeed, we could produce expression maps that were highly correlated to 20X imaging, both detecting individual spots and gene density, but with higher throughput (**Fig. 2F and Fig. S7B**). This could be an alternate strategy for imaging in a sequential manner and a good candidate for compressed sensing^(15)^.

### Comparison to published osmFISH dataset

In order to evaluate how well the unsupervised clustering works with the dRNA method, we compared the dataset acquired to that published by Codeluppi *et al.* implementing osmFISH^(4)^. We cropped our dataset to a similar region of the somatosensory cortex, CA1 of hippocampal formation, and lateral ventricle. We then re-clustered our dataset using the same 33 genes with the aim of obtaining comparable clusters. In this region, we defined 43 clusters compared to the 32 clusters found in the osmFISH dataset (**Fig. 3A and Fig. S8A**), that was spatially mapped back onto the tissue (**Fig. 3B and Fig. S8B**). Correlation between the mean expression of different clusters shows high correspondence for most of the clusters found with the two techniques (**Fig. 3C**). Overall, the agreement was best for the more divergent cell-types while for subtypes of excitatory and inhibitory neurons did not align very well, where HybRISS resolves more excitatory cell clusters. The comparison of the clusters found by the both methods and published single-cell RNA-sequencing (scRNA-seq) shows that, despite having lower detection efficiency, dRNA-HybRISS is able to define cell types with a similar resolution level as osmFISH (**Fig. S9A,B**). A more elaborate description of this comparison is presented in **Supplementary Note 2**.

**Figure 3:**
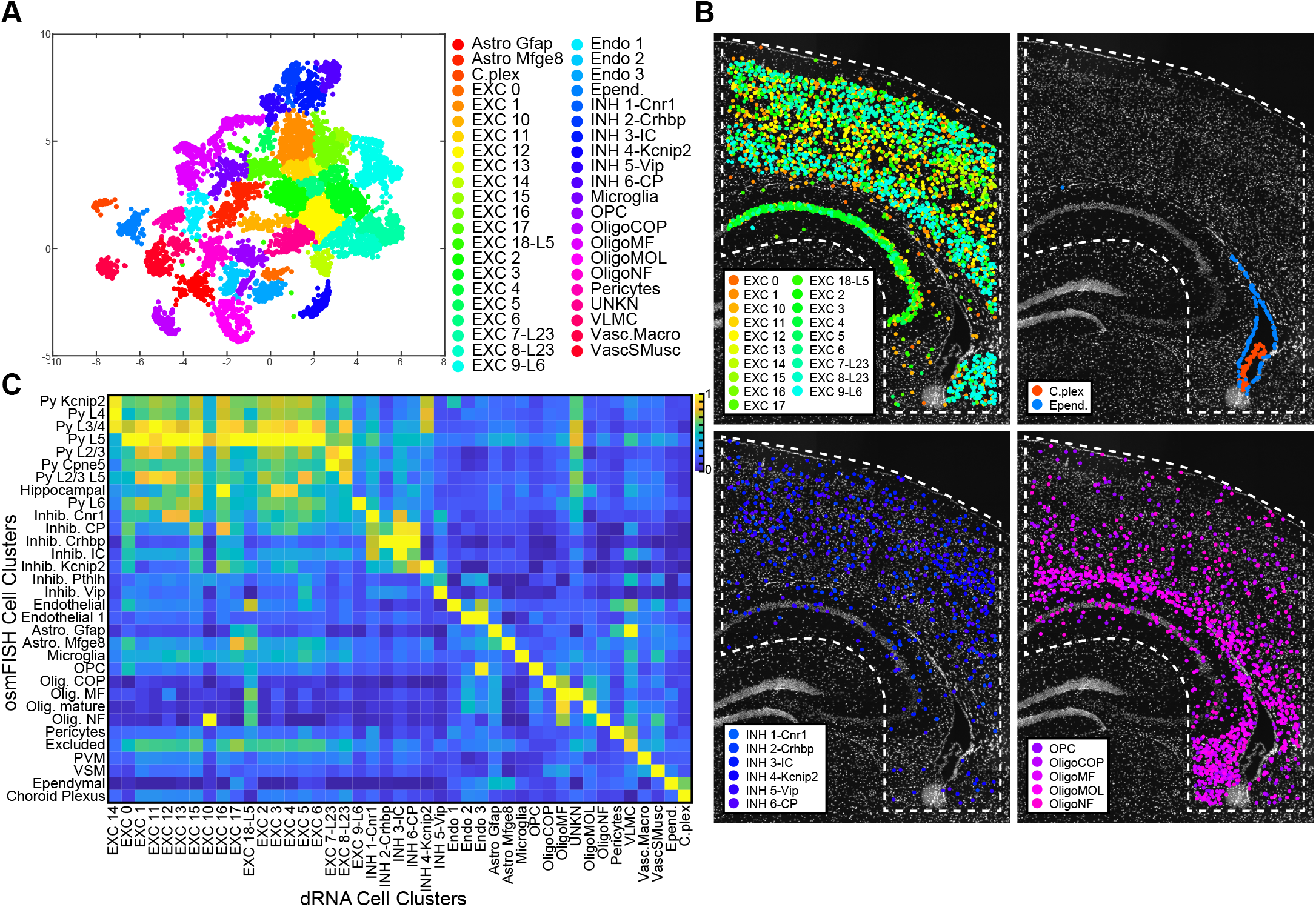
*De novo* clustering of somatosensory cortex region and comparison to osmFISH cell type clustering. (**A**) UMAP of cell clusters using 33-gene panel within outline ROI. (**B**) Cell-type map of most cell clusters superimposed on DAPI nuclear image. All cell clusters mapped in Fig. S8B. (**C**) Correlation map of osmFISH cell clusters compared to dRNA-HybRISS de novo clusters.

## DISCUSSION

The HybRISS method presents itself as an improved alternative to the traditional cDNA-based HybISS technology^(9)^ as it demonstrates a five-fold improvement in transcript detection efficiency, while maintaining specificity, and same degree of throughput and multiplexing capabilities. This increase in detection efficiency closes the gap in the analytical capabilities to that of other FISH-based techniques. Moreover, the high SNR achieved from RCA enables the use of a conventional epifluorescence microscopes, with low magnification objectives (20X-40X) at good resolution for robust spot calling and decoding with high throughput. Here we show that even lower magnification objectives (10X) can be used to identify the level of expression of each cell based on signal density and possibly in combination with compressed sensing strategies^(15)^ as a solution for optical crowding. Being able to scan large areas obtaining enough molecular information of each individual cell will be key in cell atlas projects or to support to biological questions in large tissue samples.

Currently, alternative protocols that involve the direct probing of RNA with PLPs and RCA, such as SCRINSHOT^(16)^ and targeted ExSeq^(17)^ have also exhibited improved transcript detection efficiency. However, PBCV-1 DNA ligase used in these protocols have shown high tolerance for mismatches in ligation and extensive optimization would have to be undertaken for different tissue types to prevent off-target detection^(13,16)^. The increased efficiency of dRNA has allowed for improved detection of lower expressed transcripts, which would otherwise be challenging to detect with the cDNA approach. This enables more data being generated per cell, providing opportunities to reach conclusions to a wider range of biological questions. The increased detection efficiency has potential drawback of risk to optical crowding where individual RCPs can’t be distinguished from each other using combinatorial decoding. In order not to confuse the comparison with the cDNA-based method due to crowding, we performed decoding in a non-combinatorial fashion in this work. This is a feasible approach if multiplexing levels are relatively low, such as a 50 gene panel that can be decoded in 10 rounds of 5-plex imaging per round, particularly when using 10X imaging. For higher multiplexing, one can group genes for an optimal combinatorial experiment without optical crowding by using prior knowledge from, for example, scRNA-seq data sets. Alternatively, a combination of combinatorial and non-combinatorial decoding cycles can be applied which adds on experimental and imaging rounds, but enables generation of dense, yet not optically crowded data.

As with all spatial methods, depending on the biological question being asked, the ideal method should be chosen. For example, our current cell-typing pipeline, pciSeq, does not require high detection efficiency per cell to robustly define cell types^(11)^, making the traditional cDNA-based HybISS sufficient for cell type mapping. However, should one aim to identify low abundant transcripts, or for example SNPs and mutations, dRNA-HybRISS would be a better option. Additionally, dRNA targeting would be a useful method for FFPE tissues where RNA is more degraded.

To further evaluate dRNA-HybRISS, in its multiplex capability and the data quality that it generates, we were able to cluster cell types in a mouse brain section with a panel of 50 genes and compare it to published data sets from both scRNA-seq and osmFISH. From our 50-plex experiment data, we were able to robustly decode 50 genes sequentially to confidently identify cell clusters, which have shown good correlation with both scRNA-seq and osmFISH data set, pointing to the fact that the HybISS chemistry is very much compatible with highly multiplexed experiments and also generates high quality data. The full potential of dRNA-HybRISS has yet to be explored. Nonetheless, we believe that dRNA-HybRISS can be a powerful tool for cell typing especially when combined with scRNA-seq data for gene target selection.

## METHODS

### Probe selection and design

Genes were selected based on previous publications to delineate cell types in adult mouse brain sections. Subsets of the 50-plex panel were taken from (4) and (11). The 4-plex panel is contained within the 50-plex panel. Gene lists were sent to CARTANA with accompanying customized ID sequences for in-house HybISS chemistry detection. For the 4-plex gene assay, probes were designed by CARTANA to target matching complimentary mRNA and cDNA to suit the different chemistries for benchmarking studies. Target sequences and PLP design is CARTANA proprietary information and are unknown to users and only targeted exons and number of probes used are known (**Table S1**). Mouse reference genes for cDNA-HybISS method were designed as previously published^(9)^ and sequences found in **Table S1**.

### Tissue

Mouse tissue was obtained from the Allen Brain Institute under the SpaceTx consortium. Fresh whole mouse brain tissue was cryopreserved in optimal cutting temperature (OCT) and sectioned with a cryostat (CryoStar™ NX70) at 10 μm and collected on SuperFrost Plus microscope slides. Slides stored at −80°C were air dried for five minutes and the fixed in 3% paraformaldehyde solution before respective protocols were performed.

### cDNA-HybISS protocol

The protocol was followed as published^(9)^ and at <protocols.io> (dx.doi.org/10.17504/ protocols.io.xy4fpyw). As with all dRNA probes, cDNA probes for the 4-plex assay were also provided by CARTANA to match complementary sequences of the dRNA target sequences.

### dRNA-HybRISS protocol

CARTANA provided reagents in kits (High-Sensitivity library preparation kit) with an accompanying protocol that was followed. Briefly, after tissue fixation, probe mix was incubated on tissue section overnight in hybridization buffer followed by stringent washes and then incubated in a ligation mix. After washes, RCA was performed overnight and labelled for detection. Protocols for both RCA and detection are identical to cDNA-HybISS.

### IHC staining protocol

After HybRISS RNA detection, tissue was blocked with PBTA (PBS, 5% normal donkey serum (Jackson ImmunoResearch), 0.5% Triton-X 100) for one hour. Then sections were incubated with primary antibodies, either MBP (Abcam, ab7349) or TUBB3 (BioLegend, 801213) overnight at +4°C. Sections were then washed three times with PBS and incubated with secondary antibodies (Alexa Fluor anti-rat 488 and anti-mouse 555) for 2 hours at room temperature and counterstained with DAPI.

### Imaging

All images were obtained with a Leica DMi8 epifluorescence microscope equipped with an external LED light source (Lumencor^®^ SPECTRA X light engine), automatic multi-slide stage (LMT200-HS), sCMOS camera (Leica DFC9000 GTC), and objectives (HC PL APO 10X/0.45; HC PL APO 20X/0.80; HCX PL APO 40X/1.10 W CORR). Multispectral images were captured with microscope equipped with filter cubes for 6 dye separation and an external filter wheel (DFT51011). Image scanning was performed by outlining ROIs that could be saved for multi-cycle imaging tiled imaging with 10% overlap. Z-stack imaging of 10 μm at 0.5 μm steps to cover the depth of the tissue.

### Image processing

Imaging data was processed and analyzed with an in-house pipeline based on the programming language MATLAB. All associated software can be found in a repository (https://github.com/Moldia/HybrISS).

Maximum intensity projection was performed on each field of view in order to obtain a 2-dimensional representation of each tile. Then, stitching of tiles was performed using a MATLAB implementation of MIST^(18)^ algorithm, obtaining, after exporting, different *tiff* images corresponding to each channel and round. After aligning the images and top-hat filtering them, signals were identified by manually defining an intensity and size threshold on each channel. For experiments including multiplexing, a spot-associated quality score was calculated by dividing the intensity score of the channel where the signal was detected by the sum of the intensities of all the other channels, excluding DAPI. Assuming a perfect alignment between images, each signal in the 1^st^ cycle was associated with its closer signal in the 2^nd^ cycle generating a possible barcode. A quality score was given to each union, being the distance between signals expressed in number of pixels. For each of these barcodes, a final quality score (Q) was calculated as:

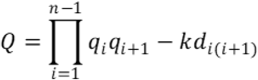

where n=2, since 2 cycles have been used in the combinatorial experiment. d_i(i+1)_ represents the distance between two signals in different rounds and is modulated by the parameter k, which can be tuned. The variables q_i_ and q_i+1_ represent the quality of a signal in the first cycle and the second respectively. Barcodes were filtered based on their final quality score (Q), keeping only those multiplexed signals presenting a high quality (Q>0).

### Cell segmentation

DAPI staining was used to identify cell nuclei by filtering its signal based on a manually set intensity threshold. Watershed segmentation was performed on top of that in order to identify approximate cell boundaries. Signals detected within the cell boundaries of a cell were assigned to it, capturing the expression profiles of individual cells.

### Data analysis

Segmented cells were filtered depending on their gene expression, selecting only cells containing more than 7 reads/cell and less than 150 reads/cell. The expression of each gene was normalized by dividing the number of reads by the standard deviation of each gene’s expression. Then, density-based spatial (DBSCAN) clustering was performed on the normalized gene/cell matrix and resulting clusters were manually annotated.

## DATA AVAILABILITY

Pre-processed images or raw tile images (several terabytes) are available from the corresponding author upon request.

## CODE AVAILABILITY

All code is available online at https://github.com/Moldia/HybrISS.

## ACKNOWLEDGEMENTS

This work was supported by Eurostars Project; The Chan Zuckerberg Initiative, an advised fund of Silicon Valley Community Foundation; Swedish Brain Foundation (Hjärnfonden) [PS2018-0012]; EASI Genomics (H2020); Vetenskapsrådet; Knut and Alice Wallenberg Foundation; Erling Persson Family Foundation. We thank Gonçalo Castelo-Branco and Ernest Arenas for donation of primary and secondary antibodies and SpaceTx consortium for tissue. Imaging performed on Leica DMi8 is thanks to Jens Hjerling-Leffler with support from ERC grant (#819540). We thank all members of the Mats Nilsson lab for their insight and comments.

## AUTHOR CONTRIBUTIONS

HL performed all experiments and analyzed data. SMS and DG analyzed data. HL, DG and MN conceived the study. DG and MN supervised the study. All authors contributed to the writing of manuscript.

## CONFLICT OF INTEREST

MN is co-founder of the company CARTANA AB from which reagent kits were obtained for this research study. All reagents used from CARTANA are listed in the Methods section.

## SUPPLEMENTARY INFORMATION

Supplementary information contains 2 Notes, 9 Figures, and 2 Tables.

## Notes

### Competing Interest Statement

Mats Nilsson is co-founder of the company CARTANA which is commercializing in situ sequencing reagents. Cartana has been acquired by 10X Genomics and Mats Nilsson currently serves as a scientific advisor to 10X Genomics.

