## Supplementary_Information for "Direct RNA targeted transcriptomic profiling in tissue using Hybridization-based RNA In Situ Sequencing (HybRISS)"

*Supplementary information contains 2 Notes, 9 Figures and 2 Tables*

**DATA**

*Supplemental Note 1: description of the clusters identified in Figure 2*
*Supplemental Note 2: comparison of clusters*

*Supplementary Fig. 1: HybISS method and decoding chemistry*
*Supplementary Fig. 2: dRNA-HybRISS method and comparison to cDNA-HybISS using 4-plex gene panel*
*Supplementary Fig. 3: Validation of dRNA-HybRISS specificity using 4-plex gene panel*
*Supplementary Fig. 4: Control experiments for dRNA-HybRISS specificity using 4-plex gene panel*
*Supplementary Fig. 5: Multiplex implementation of dRNA-HybRISS to target panel of 50 genes*
*Supplementary Fig. 6: 50 gene targeting with dRNA-HybRISS*
*Supplementary Fig. 7: 10X imaging analysis of Mbp/Lamp5/Cd24a*
*Supplementary Fig. 8: Subset gene panel and region clustering and comparison to osmFISH*
*Supplementary Fig. 9: Method comparison to single-cell RNA-sequencing defined clusters*

*Supplementary Table 1: Padlock probes and relevant sequences, bridge probe sequences, detection oligonucleotide sequences and fluorophores, and 50-plex gene imaging rounds*
*Supplementary Table 2: 50-plex individual cell expression and 50-plex main cluster expression*

#### Supplementary Note 1: Description of the clusters identified in Figure 2

A total of 56 clusters were defined after clustering the segmented data with a DBSCAN algorithm. Those clusters were annotated based on known cell type markers expressed in different clusters and the direct comparison to cell types described by Zeisel *et al.* scRNA-seq dataset of the mouse central nervous system[19] to be present in the same brain region characterized in this experiment.

A total of 20 clusters were classified as excitatory neurons, characterized by the expression of markers such as *Slc17a7* or *Calb2*. Excitatory neurons were mostly found within the cortex, the hypothalamus and, in less abundance, the thalamus. Some distinct excitatory cell types could be distinguished, such as *Excitatory 19 Car3+*, which showed a specific location in the intermediate layers of the cortex (L3-4). Eight different clusters were described to be capturing different inhibitory cell types. Two main groups could be described among the inhibitory clusters: Five clusters which cells mainly located in the cortex, and other three region-specific clusters, including two clusters located in the caudoputamen (*INH 10 Penk* and *INH Penk*) and one cluster in the reticular nucleus of the thalamus (*INH 2-Pvalb 1*). These three clusters, despite their distinct location in the UMAP (Fig. 2B) have been assigned to inhibitory clusters due to the resemblance to the inhibitory class, although they might need to be classify in a distinct group of cells.

We described the remaining 26 clusters as non-neuronal clusters, since their expression profile presents specific markers for typical non-neuronal cell types. Among these, we found 11 oligodendrocyte-like clusters, characterized by the expression of markers like *Plp1*, *Mbp* or *Sox10*. Most of the cells assigned to these clusters are found in the fiber tracts. Within these clusters, deeper classification was possible for specific clusters. Therefore, an oligodendrocyte precursor cells (OPC) cluster was described, thanks to the expression of markers such as *Pdgfra*, showing a broader location than the rest of oligodendrocytes, with presence of some cells in the cortex and thalamus. Additionally, clusters matched with committed oligodendrocyte precursors (COP), myelin-forming oligodendrocytes (MFOL) and newly formed oligodendrocytes (NFOL) cell types.

We described two different clusters showing an astrocyte-like expression profile, differentiating between *Mgfe8*$^+$ cells and *Gfap*$^+$ cells, together with a cluster with a clear microglia-like expression profile. Within non-neuronal clusters, we also defined clusters assigned to microglia (1), Smooth Muscle cells (SMC), Pericytes (3), endothelial cells (1), vascular and leptomeningeal cells (VLMC) (1), vascular endothelial cells (1), ependymal cells (1) and choroid plexus (1).

Some other clusters didn't match well with any specific cell type described in Zeisel *et al.*[19], but were characterized by the expression of certain genes, showing a clear location in the tissue. This is the case of *Tac2*$^+$ cells, located in the medial habenula and *Calb2*$^+$ cells, placed in specific regions of the thalamus. In the case of *Aldoc*$^+$ cells, no specific location was found, but cells were characterized by a high expression of *Aldoc*. Finally, two different clusters could not be annotated, since they present a wide expression across the tissue and their expression profile didn't match any specific cell type, having have levels of expression for both neuronal and non-neuronal markers. We believe these clusters are artifacts in part due to a bad segmentation of certain cells across the tissue where close neighboring cells are segmented as one.

#### Supplementary Note 2: Comparison of clusters

In order to compare the expression of the clusters defined by dRNA-HybRISS and by osmFISH using the list of 33 genes used in Codeluppi *et al.*[4] and the cell types defined using scRNA-seq data from Zeisel *et al.*[19], the correlation between the mean expression of each cluster on each method was calculated (Fig. 3C).

Correlation between the mean expression of different clusters shows high correspondence for most of the clusters found with the two spatial techniques, even though dRNA-HybRISS presents a much lower detection efficiency. One-to-one correspondence between clusters was observed for many of the of the non-neuronal cell types, including endothelial cells, perivascular macrophages (PVM), vascular smooth muscle cells (VSM), choroid plexus, microglia, astrocytes, pericytes and ependymal cells. Regarding clusters expressing oligodendrocyte markers, we were able to clearly distinguish oligodendrocyte precursor cells (OPC) and partially distinguish committed oligodendrocyte precursor cells (Olig. COP), but further subtypes were not clearly distinguishable.

Within neuronal clusters, some discrepancies were present between the inhibitory neuron subtypes. The most important discrepancies between the methods were found among the excitatory cells where both methods showed poor capacities to resolve cell types clearly. In this case, no clear one-to-one correspondence can be found within excitatory clusters, except for Pyramidal L4 and Pyramidal L6, where a correlation between a specific osmFISH and a dRNA-HybRISS cluster is observed.

Our clusters were also compared with the cell types described by Zeisel *et al.*[19] by scRNA-seq (Fig. S9A). As observed when comparing the dRNA clusters with the osmFISH ones, clear correspondence between non-neuronal clusters and some distinct neuronal populations was found. However, one-to-one correspondence between most excitatory, as well as some inhibitory and oligodendrocyte cell types was not achieved. Similar conclusions are extracted when comparing the osmFISH dataset with the scRNA-seq based cell types, where osmFISH clusters do not match perfectly the scRNA-seq ones for a considerable amount of excitatory and inhibitory populations (Fig. S9A).

The comparison of the clusters found by the three methods shows that, despite having lower detection efficiency, dRNA-HybRISS is able to define cell types with a similar resolution level as osmFISH. Discrepancies between the clusters defined by scRNA-seq clustering and both spatial methods are consistent, proving the importance of the gene panel curation in targeted methods like these.

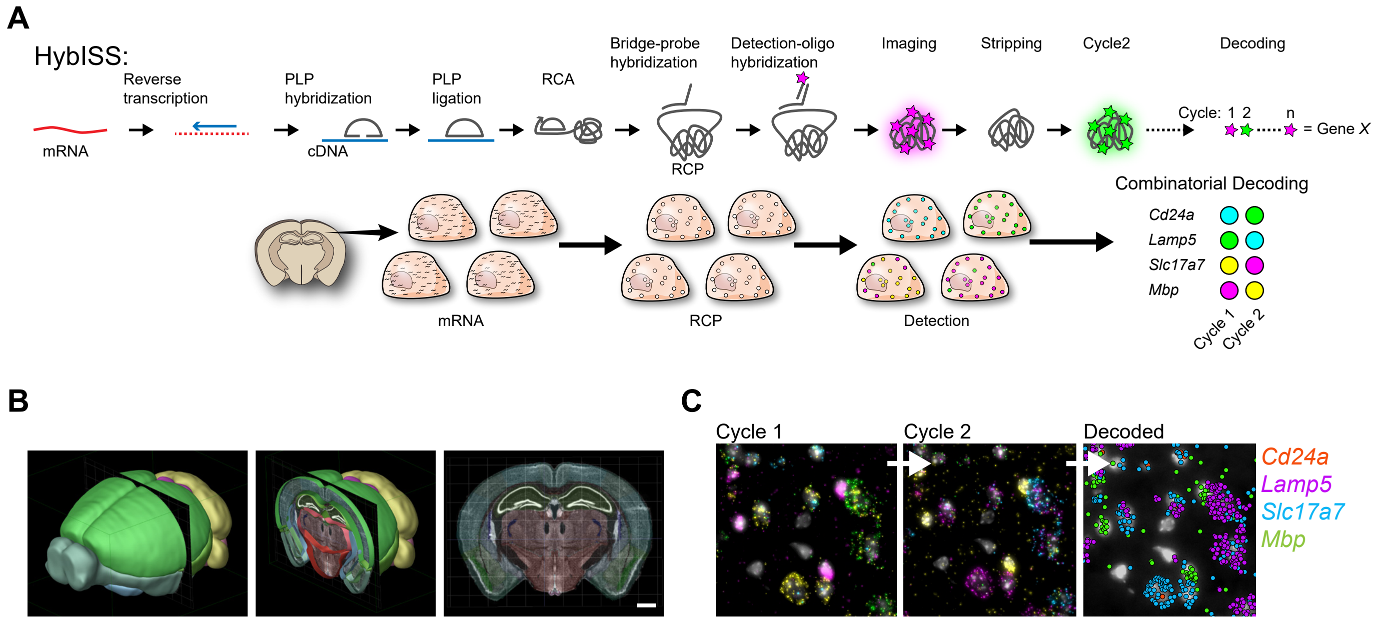


**Supplementary Figure 1: HybISS method and decoding chemistry.**

*(**A**) Schematic of HybISS chemistry: reverse transcription for cDNA synthesis, PLP hybridization and ligation, RCA, RCP detection, imaging, stripping and repeated cycles for combinatorial barcode decoding.*

*(**B**) Regional localization of mouse brain coronal section used for analysis. Image credit: Allen Brain Institute.*

*(**C**) Example of combinatorial decoding that is possible with HybRISS and gene panels used.*

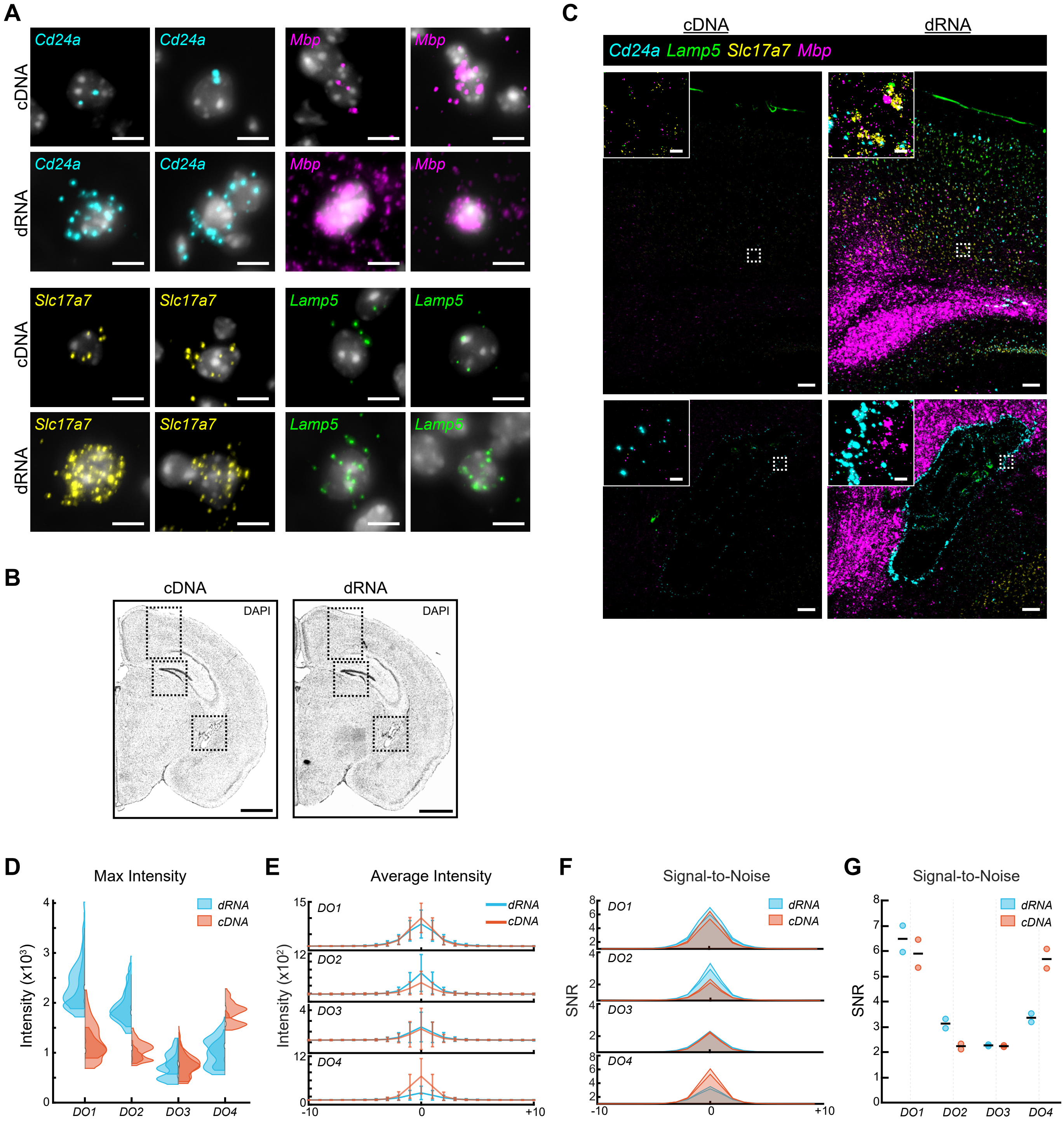


**Supplementary Figure 2: dRNA-HybRISS method and comparison to cDNA-HybISS using 4-plex gene panel.**

*(**A**) Representative images of single cells showing increased detection of individual transcripts in cDNA-HybISS and dRNA-HybRISS. Scale bar, 5 µm.*

*(**B**) DAPI image of half coronal sections for cDNA and dRNA with demarked ROI regions used for analysis.*

*(**C**) Representative raw images from two of the three ROIs (B) and the distribution of the 4-plex panel. Experiments run in parallel and same postprocessing intensitysignal level adjustments. ROIs include regions of cortex (top) and lateral ventricle (bottom). Scale bar, 100 µm, inset 10 µm.*

*(**D**) Violin plots displaying max intensity of top 50 RCPs in three ROIs from two samples in each channel measured for cDNA-HybISS and dRNA-HybRISS. DO1-AF750, DO2-AF488, DO3-Cy3, DO4-Cy5.*

*(**E**) Average intensities of measured RCPs in all ROIs (three from each of the two samples for each condition). Pixel intensity measured across a 21-pixel line bisecting RCPs.*

*(**F**) SNR measured from (E). Outer 2-pixel measurements at each end of 21-pixel line were used to calculate background noise. Each pixel intensity measurement was divided by the noise to display SNR across the RCPs.*

*(**G**) Peak SNR from (F) measured across four fluorescent channels.*

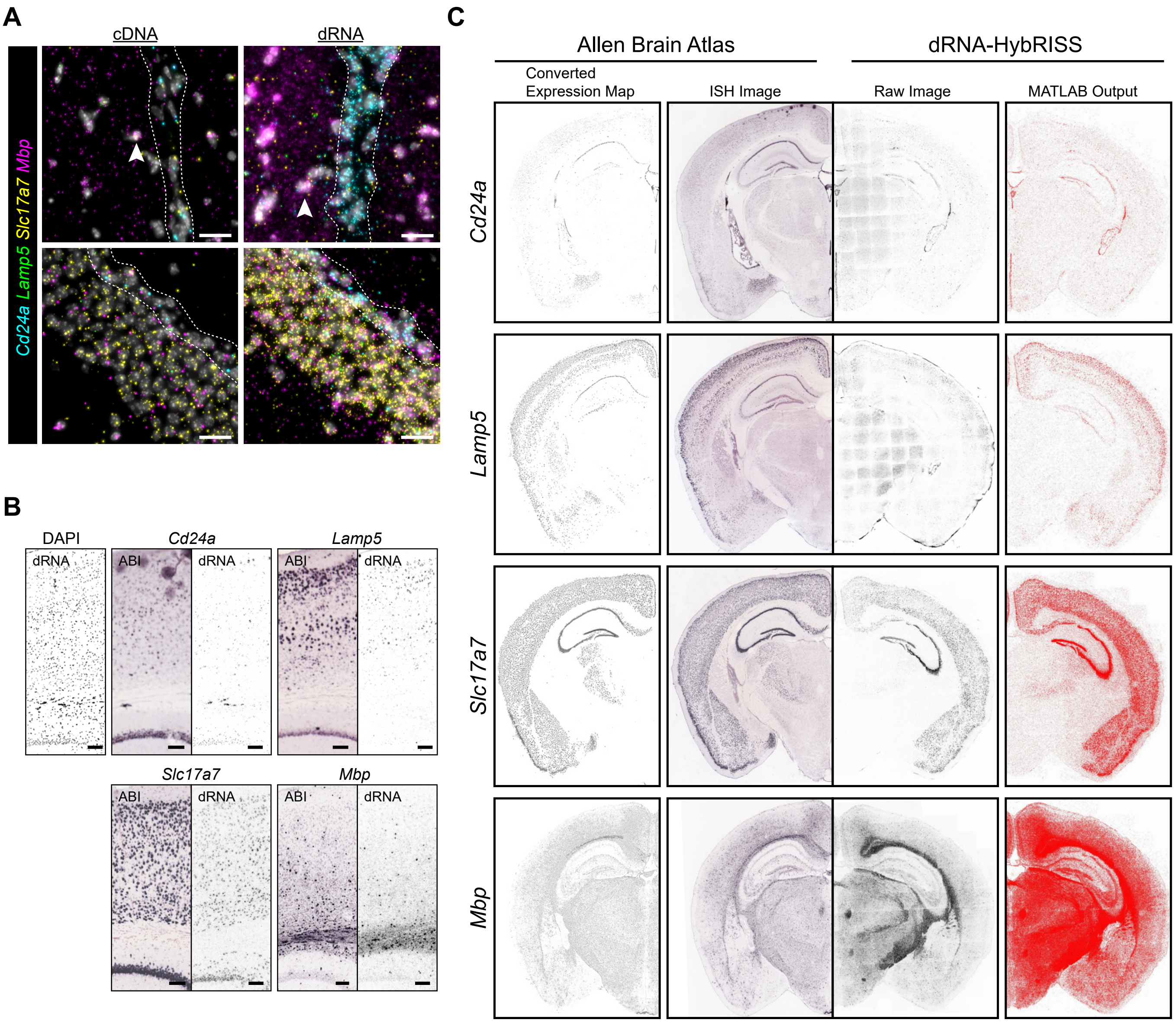


**Supplementary Figure 3: Validation of dRNA-HybRISS specificity using 4-plex gene panel.**

*(**A**) Distribution of 4-plex genes around ventricle (top) and hippocampus (bottom). Scale bar, 20 µm.*

*(**B**) Transcript distribution of 4-plex panel compared to Allen Mouse Brain Atlas (ABI) in a region of the cortex. Image credit: Allen Institute.*

*(**C**) Transcript distribution of Cd24a/Lamp5/Slc17a7/Mbp compared to Allen Mouse Brain Atlas[20] in half coronal section. Image credit: Allen Institute.*

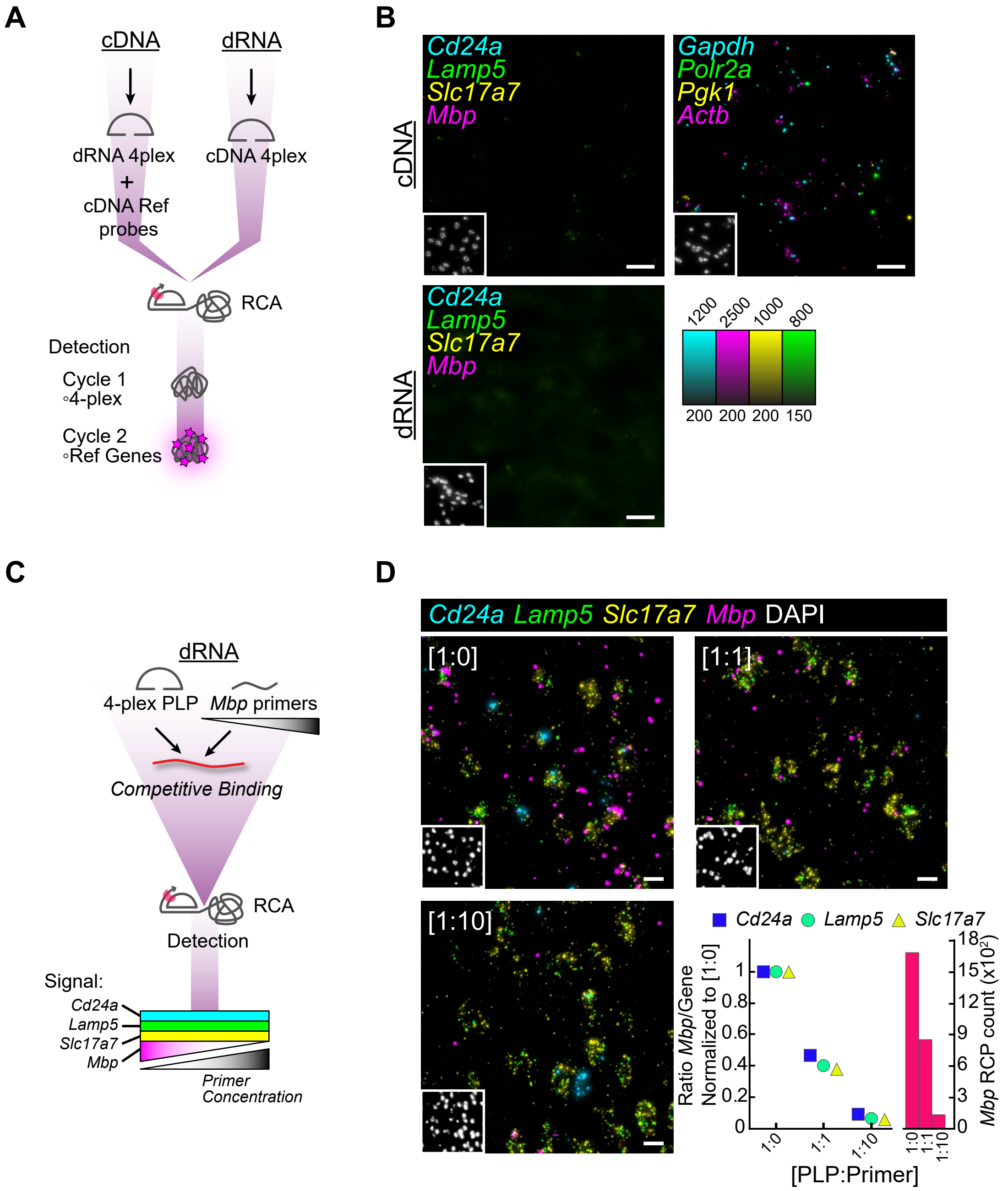


**Supplementary Figure 4: Control experiments for dRNA-HybRISS specificity using 4-plex gene panel.**

*(**A**) Schematic of negative control experiment where probes for cDNA and dRNA protocols were swapped in each respective protocol. A panel of mouse reference cDNA probes were added into the probe pool for a positive control for cDNA protocol and detected in the second round.*

*(**B**) Raw images of experiment in (d) indicating specificity of probes. Inset, DAPI. Scale bar, 10 µm.*

*(**C**) Experimental schematic of competitive assay with different concentrations of primers that bind to Mbp target sequences, and Cd24a/Slc17a7/Lamp5 as controls.*

*(**D**) Images from competitive assay and calculation showing ratio of Mbp to all other genes and total count of Mbp.*

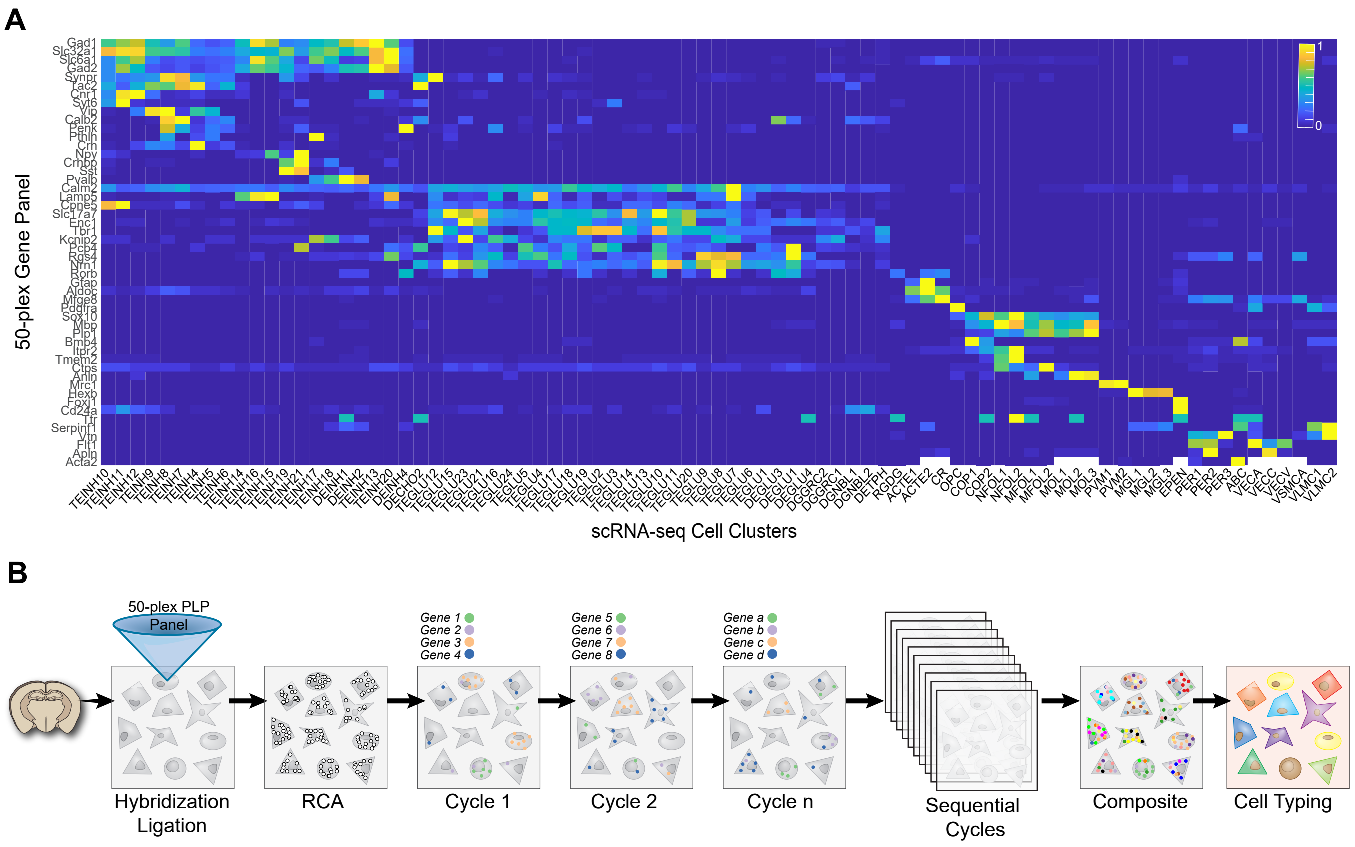


**Supplementary Figure 5: Multiplex implementation of dRNA-HybRISS to target panel of 50 genes.**

*(**A**) 50-plex gene selection representation in Zeisel et al. scRNA-seq data[(19)].*

*(**B**) Schematic overview of experimental workflow of 50-plex gene panel and sequential decoding in order to produce a composite image and cell typing based on transcript expression.*

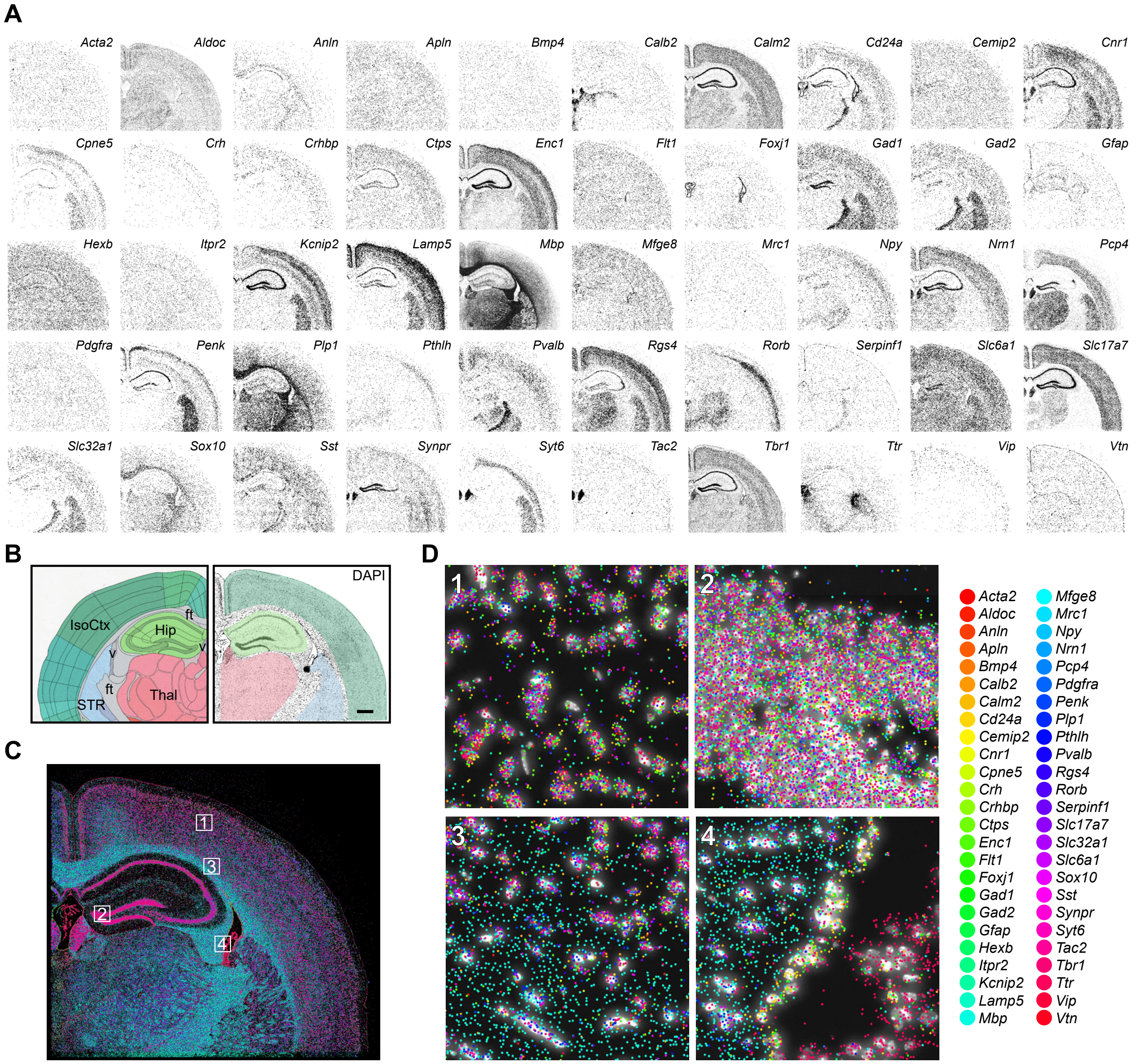


#### Supplementary Figure 6: 50 gene targeting with dRNA-HybRISS.

(**A**) *Decoded images of all genes in the 50-plex panel imaged over 14 rounds in region of coronal mouse brain.*

(**B**) *Anatomical position of mouse coronal brain section used in 50-plex experiment (right, area of 0.216 mm²) and Allen Mouse Brain Reference Atlas (left) covering isocortex, hippocampus, thalamus, striatum, ventricles, and fiber tracts. Scale bar, 500 µm.*

(**C**) *50-plex gene panel, 14 round composite images in coronal mouse brain section.*

(**D**) *Regions of interest from (C) showing the transcript density and diversity.*

### Supplementary Figure 7

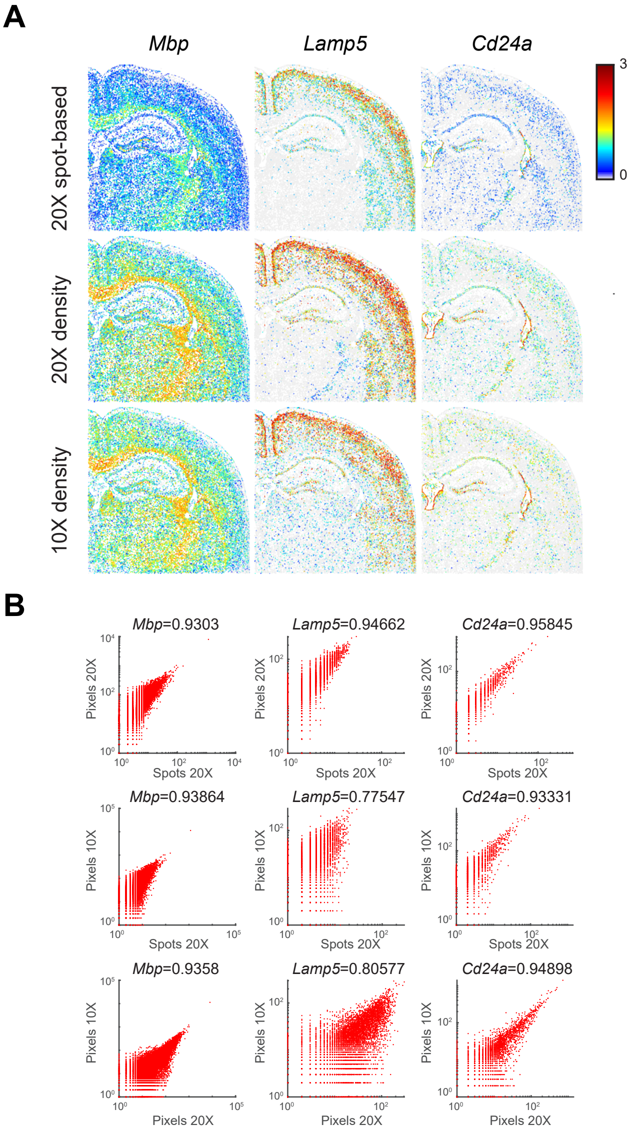


**Supplementary Figure 7: 10X imaging analysis of *Mbp/Lamp5/Cd24a*.**

*(**A**) 20X objective spot-based detection along with 20X density-based detection and how it compares to 10X objective density-based transcript detection of Mbp, Lamp5, Cd24a.*

*(**B**) Expression correlation of pixel and spot detection for 20X objective and same comparison between 10X pixels and 20X spot detection.*

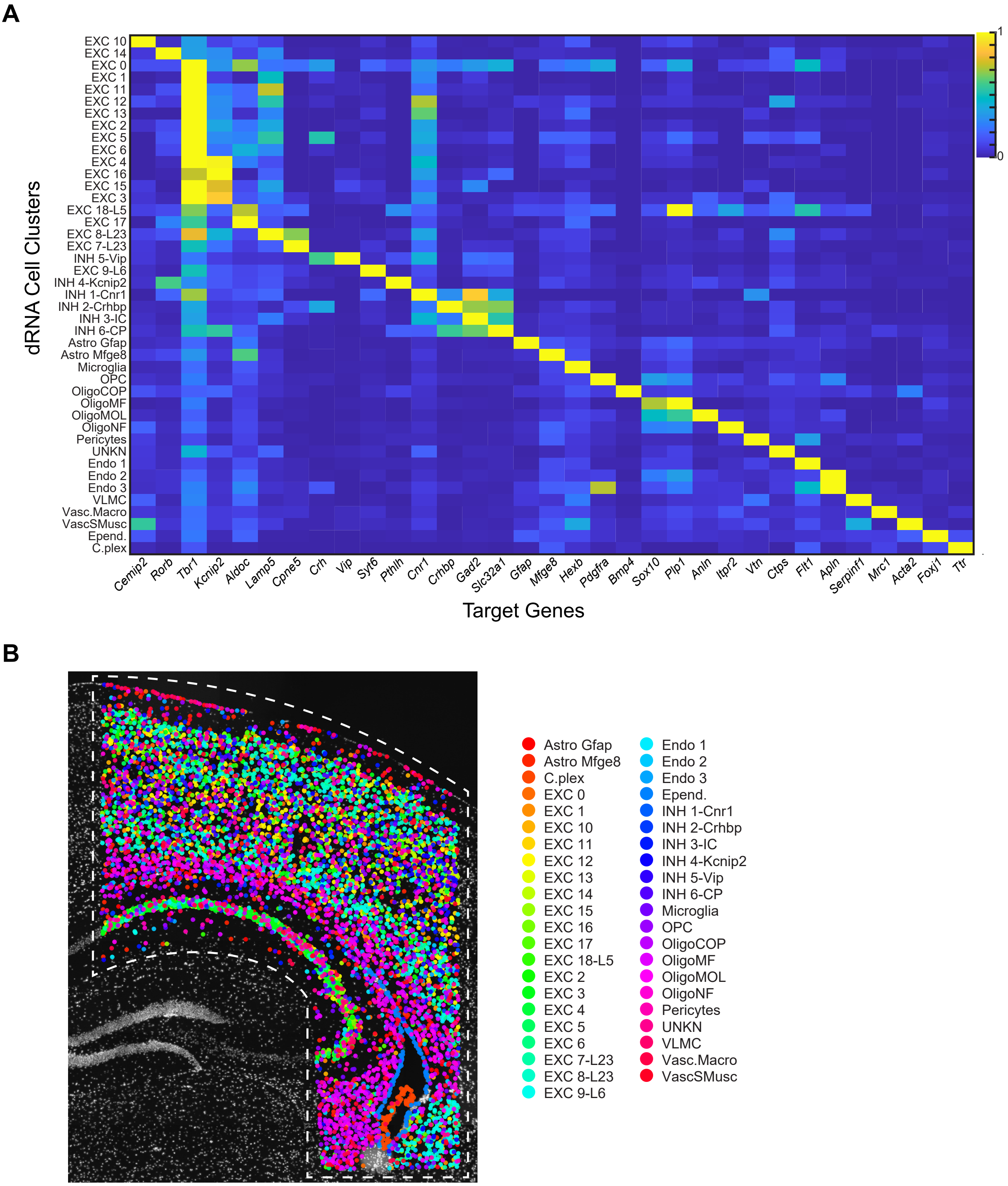


**Supplementary Figure 8: Subset gene panel and region clustering and comparison to osmFISH.**

*(**A**) De novo clustering of spatial data based on segmented ROI and 33-gene subset taken from the osmFISH panel[(4)].*

*(**B**) Expression map of all cell clusters superimposed on DAPI nuclear image.*

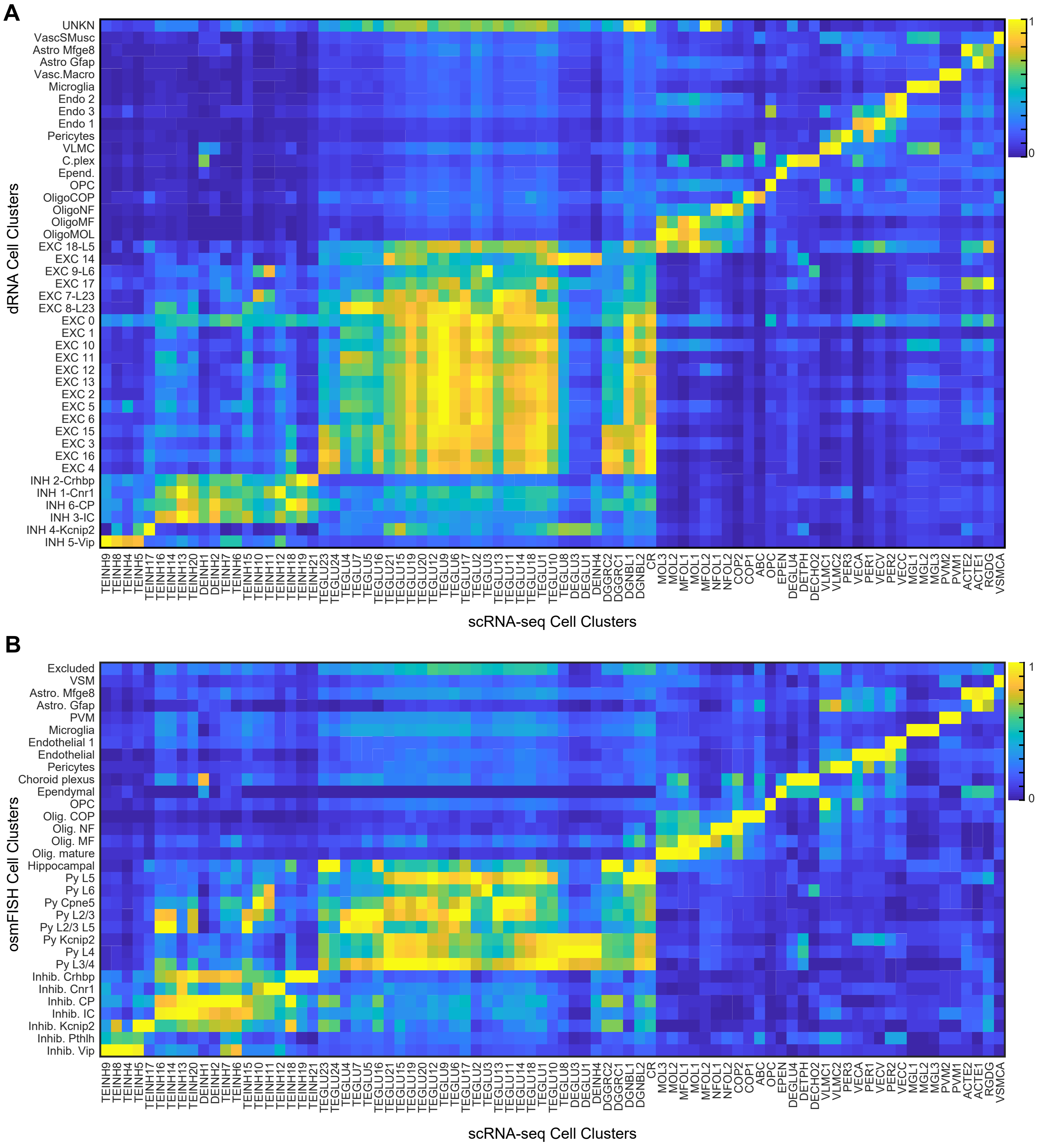


**Supplementary Figure 9: Method comparison to single-cell RNA-sequencing defined clusters.**

*(**A**) dRNA de novo clusters compared to scRNA-seq clusters[(19)].*

*(**B**) osmFISH cell clusters compared to scRNA-seq data[(19)].*
